# A direct RNA-seq-based EBV Latency Transcriptome Offers Insights into the Biogenesis of EBV Gene Products

**DOI:** 10.1101/2025.04.23.648900

**Authors:** Aaron Mamane-Logsdon, Isabelle Zane, June See Chong, Oscar Hou In Chou, Jiajun Huang, Mahesh Rawal, Adam C. Gillman, Wiyada Wongwiwat, Mostafa Saleban, I’ah Donovan-Banfield, David A. Matthews, Robert E. White

## Abstract

Epstein-Barr virus (EBV) ubiquitously infects humans, establishing lifelong persistence in B cells. In vitro, EBV-infected B cells can establish a lymphoblastoid cell line (LCL). EBV’s transcripts in LCLs (Latency III) produce six nuclear proteins (EBNAs), two latency membrane proteins (LMPs) and various microRNAs and putative long non-coding RNAs (BARTs). The BART and EBNA transcription units are characterised by extensive alternative splicing.

We generated LCLs with B95-8 EBV-BACs, including one engineered with “barcodes” in the first and last repeat of internal repeat 1 (IR1), and analysed their EBV transcriptomes using long-read nanopore direct RNA-seq. Our pipeline ensures appropriate mapping of the W promoter (Wp) 5’ exon, and corrects W1-W2 exon counts that misalign to IR1. This suggests that splicing across IR1 largely includes all W exons, and that Wp-derived transcripts more frequently encode the EBNA-LP start codon than Cp transcripts. Analysis identified a short variant of exon W2 and a novel polyA site before EBNA2, provided insights into BHRF1 miRNA processing and suggested co-ordination between polyA and splice site usage, although improved read depth and integrity are required to confirm this. The BAC region disrupts the integrity of BART transcripts through premature polyadenylation and cryptic splice sites in the hygromycin expression cassette. Finally, a few transcripts extended across established gene boundaries, running from EBNA, to BART to LMP2 gene regions, sometimes including novel exons between EBNA1 and the BART promoter. We have produced an EBV annotation based on these findings to help others better characterise EBV transcriptomes in future.

**Data Summary:** Scripts (and the shell scripts used to combine commands into the pipeline), and guidance in their usage are available on Github (github.com/robertewhite/ebv-transcriptomics-tools). All of the RNA-seq raw data and analyses relevant to this study are available from the EMBL Nucleotide Archive Study accession PRJEB83447 (read files ERR14129300-303), or from the author’s website, ebv.org.uk. Processed and analysed data is presented in Excel format in Supplementary tables ST1-8, and the intermediate data processing conducted in excel is linked from the front page of ebv.org.uk, alongside the scripts, raw reads and updated B95-8-BAC and prototype EBV transcriptome annotation (gff3) files.

**Impact statement:** This article showcases the potential of direct RNA-seq to characterise complex transcriptomes like Epstein-Barr virus, highlights specific challenges of interpreting RNA-seq data, and presents tools to solve specific challenges of mapping RNA-seq reads to EBV exons. Biologically, the data analysis identifies several new aspects of Epstein-Barr virus transcription, offering insights into miRNA proteins, and identifies idiosyncrasies of the transcriptome of the widely used B95-8 BAC that will inform studies using this system. Finally we provide an improved annotation for EBV RNA-seq studies.

## Introduction

Epstein-Barr virus (EBV) is a ubiquitous human gamma-herpesvirus, that establishes life-long persistence in resting memory B cells in the vast majority of the adult population. EBV establishes a persistent latent state, transitioning to productive replication when these differentiate into plasma cells, or when infecting mucosal epithelia, where it is shed into body fluids such as saliva, which facilitates transmission. Immunodeficiency and/or mutations that disrupt normal virus-host relationships in these cell types can promote the wide range of B cell and epithelial malignancies in which EBV is implicated.

Infection of B cells in vitro results in the activation and hyper-proliferation of B cells, that can lead to the outgrowth of transformed lymphoblastoid cell lines (LCLs). Different virus strains have different transformation efficiencies, with the lab strain B95-8 being widely used due to its ready ability to generate LCLs. This strain of EBV was cloned in an F-factor plasmid (better known as a bacterial artificial chromosome, or BAC) 25 years ago [1], enabling the genetic modification of the EBV genome in bacteria, as a reverse genetic system for studying EBV gene products.

All herpesviruses are large DNA viruses, that generate a large number of transcripts, many of which overlap, with shared polyadenylation sites, which makes it challenging to accurately quantitate transcript abundance from short-read RNA-seq data. Nevertheless, considerable progress has been made in characterising the transcriptome of the productive cycle of EBV [2]. However, EBV has a particularly complex latency transcriptome, combining alternative splicing, polyadenylation and promoter usage to encode a diverse array of proteins and non-coding RNAs. The transcription programme found in LCLs (termed Latency III) has three main zones of transcription: The EBV nuclear antigen (EBNA) transcripts; the latency membrane protein (LMP) transcripts; and the BamHI A rightward transcripts (BARTs), shown schematically in **Figure 1**.

**Figure 1.**
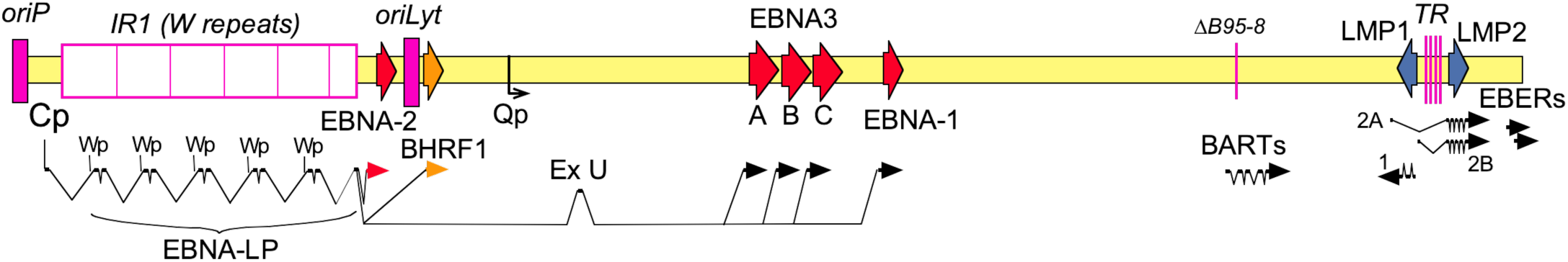
Schematic overview of main latency transcripts of B95-8 EBV. Gene and repeat structure of the virus genome (normally circular, but viewed with its ends at oriP) is shown above an overview of the main transcript structures. Gene and transcript arrows indicate transcription direction. Genes are colour-coded as EBV nuclear antigens (EBNA - red) and latency membrane proteins (LMPs – blue). EBNA promoters (Cp, Wp and Qp) and the U exon are labelled. Repeat regions (magenta) are family of repeats in oriP, internal repeats IR1 (Bam W repeat); IR2 (part of oriLyt); IR3 (glycine-alanine repeats in EBNA1), while B95-8 deletion (ΔB95-8) loses IR4 and contains the BAC (not shown); Terminal repeats (TR) lie between LMPs. Note that some genes have more than the indicated number of splicing events.

The EBNA transcripts are the first to be transcribed after infection. In genomic order: EBNA-LP (EBNA-leader protein) facilitates expression of the other virus genes by preventing genome silencing by antiviral proteins [3, 4]; EBNA2 initiates the activation and proliferation of the B cells [5], while the EBNA3 proteins induce epigenetically stable changes to gene expression that prevent apoptosis and facilitate germinal centre differentiation [6]. Finally, EBNA1 – through binding to the EBV latency replication origin, oriP – both facilitates viral gene expression and mediates the replication and segregation of the extra-chromosomal EBV genome in synchrony with the cell cycle.

EBV transcription begins using a promoter (Wp) within the major internal repeats (IR1, or BamW repeats): there is a copy of Wp in each of the 3kb repeat units of IR1. Then, once newly produced EBNA1 has established oriP [7], and EBNA2 has bound to Cp [8] and to regulatory regions upstream of the first Wp [9], transcription largely switches from Wp to the BamC promoter (Cp) that lies just upstream of IR1. Both Cp and Wp-derived transcripts are alternatively spliced and polyadenylated to variously encode the six EBNA proteins and the pro-survival Bcl2 homologue, BHRF1 [10, 11]. In addition, three miRNAs (mirBHRF1-1,-2 and -3) are processed from sites either side of the BHRF1 ORF [12, 13], while some RNAs excised from pre-mRNAs are stable non-coding RNAs of unknown function [14, 15]. In vivo, as EBV progresses to its final latency state, Cp and Wp shut down, switching to Qp that makes only EBNA1 transcripts.

On the other side of oriP from Cp are two genes for EBV-encoded RNAs (EBERs), encoding abundant RNA polymerase III-transcribed non-coding RNAs around 170 nucleotides in length [16, 17]. Slightly further from oriP, and spanning the viral terminal repeats, lie the latency membrane protein genes LMP1 and LMP2. The LMP1 and LMP2A proteins provide signals required for B cell survival during affinity maturation, while LMP2B lacks the LMP2A signalling domain through alternative promoter usage: A divergent promoter region gives rise to LMP1 and LMP2B transcripts [18], with the LMP2A promoter situated beyond the LMP1 pA site, regulated by a CTCF-binding region between LMP2A exon and LMP1 [19].

Finally, the Bam A rightward transcripts (BARTs) are a collection of alternatively spliced RNAs that span an approximately 20 kb region of the EBV genome [20]. The BART RNAs appear to be nuclear long non-coding RNAs [21, 22], but some have the potential to produce proteins (A73; RPMS1; RK-BARF0), although these proteins have yet to be detected in EBV-infected cells [21]. The BART introns produce around 40 miRNAs from 22 pre-miRNA stem-loops, mainly in two clusters [23, 24], that modulate a wide range of cell functions [25, 26]. However, much of the BART region is deleted in the commonly used B95-8 strain, and the B95-8 reverse genetic system has a bacterial artificial chromosome (BAC) and expression cassettes for GFP and hygromycin phosphotransferase inserted into this deletion, the impact of which is unknown.

The complexity of the splicing in EBV makes the quantitation of distinct transcripts from RNA-seq data challenging, as simplifications of the transcriptome annotation may offer misleading indications of the abundance of different proteins. Therefore we aimed to develop a pipeline to accurately determine and quantify EBV transcript diversity, and assess any co-dependencies of alternative splicing options in the EBV latency transcriptome of the B95-8 BAC, using long-read direct RNA sequencing (dRNA-seq) developed by Oxford Nanopore. Directly sequencing the RNA molecules avoids amplification biases while potentially reading intact whole transcript molecules. This comes at the cost of base-calling accuracy, which is less developed for RNA than DNA, and complicated by the larger range of RNA modifications [27, 28]. Lower nucleotide accuracy is mitigated by the ability to sequence whole transcripts (except for 10-12 nucleotides at the 5’ end) in a single read, allowing quantification of different combinations of splice sites, transcription start sites and polyadenylation sites of individual reads, even across repetitive regions of the genome like IR1.

Here we present a dRNA-seq pipeline with EBV-specific modifications (to cope with the small 5’ exon of Wp, and the repetitiveness of IR1 exons) to characterise the latency transcriptome of LCLs established with BAC-cloned B95-8. We demonstrate how this allows characterisation and quantitation of EBV’s most complex transcripts, and provide a detailed analysis of the B95-8 EBV-BAC’s transcription and splicing profile in LCLs.

## Results

### Generation of a high quality long read transcriptome for EBV

Our study analyses transcripts from four LCLs from three distinct variants of the B95-8 strain EBV BAC. First our clone of the original B95-8 BAC [1], which – like the parent B95-8 virus – harbours a STOP codon at the end of one of its six W1 exons [29]). Second, its derivative WT^w^ whose homogeneous IR1 lacks this W1 STOP codon [3]. And third, to enable us to specifically characterise splicing of EBNA transcripts spanning IR1 exons, we generated a recombinant EBV genome containing synonymous SNPs (“barcodes”) in the first and last W2 exons of IR1 (**Supporting Figure S1**) herein designated EBV-BC. This allow quantitation of how consistently the first and last W2 exon are included in such transcripts, and was used to generate two LCLs (EBV-BC^A^ and EBV-BC^M^). All these BAC clones contain 6.6 repeat units in IR1 (i.e. 6 pairs of W1 and W2 exons).

RNA from these LCLs was directly sequenced using an Oxford Nanopore MinION v9 flow cell, reads were base-called, and aligned to the EBV genome (to both the generic WT^w^ sequence, and the matched EBV-BC or WT^HB9^ genomes – “self”) in a splice-aware manner to avoid spurious splice junctions. The origin of WT^HB9^ was set within oriP (NC_007605 position 8315) to ensure all reads could align in full, as this was the only part of the EBV genome not crossed by transcripts (and aligners cannot use circular templates). A script was developed to correct for the failure of minimap2 to align 5’ ends of reads to exon W0. A second script was then used to identify commonly seen 5’ and 3’ ends (at least 5 reads across the four samples that start within a 5 nt window), and define reads with 5’ and 3’ ends within 20 nt either side of these positions as “full length”.

Across the 4 LCLs, 3693 reads aligned to the EBV genome. Almost half of these were from the WT^w^-LCL, and across all four samples, the vast majority of reads mapped to latency loci (LMPs, EBNAs and BARTs) and GFP transgene. A few reads were classical ‘lytic’ transcripts, but their numbers were too few for any meaningful analysis in this study. There were also no EBER transcripts identified by our pipeline, most likely because RNA polymerase III transcripts lack the polyA sequence required by the nanopore library preparation.

### EBNA transcripts primarily use Cp, but Wp contributes significantly to EBNA- LP production

As illustrated in **Figure 2A**, the EBNA and BHRF1 loci are transcribed from either Cp or Wp, and the splice junction from promoter exons (W0 or C2) to the first IR1 exon (W1) has both alternative donor exons (W0, C2 or C2’ [30]) and alternative acceptors (W1 or W1’). Splicing of C2 or W0 to W1’ produces the AUG required for translation of EBNA-LP. Other splice junctions allow translation of downstream proteins [30]. Among reads that have both conventional 5’ Cp or Wp exons and a 3’ Y exon, 90% arise from Cp, and 10% from Wp (**Figure 2B**). Cp transcripts, we observed use of both C2 and C2’ exons (C2’ being found in 35-42% of Cp reads, consistent with the ≈35% in a PacBio RNA-seq of a B95-8 LCL [30]), while a very small number (≈1%) of Cp reads skipped exon C2. When the splice donor is W0 or C2’, there is about a 60:40 ratio of W1:W1’ splice acceptor usage. In contrast the C2 splice donor has a lower preference for the W1’ splice acceptor - approximately 75:25%.

**Figure 2.**
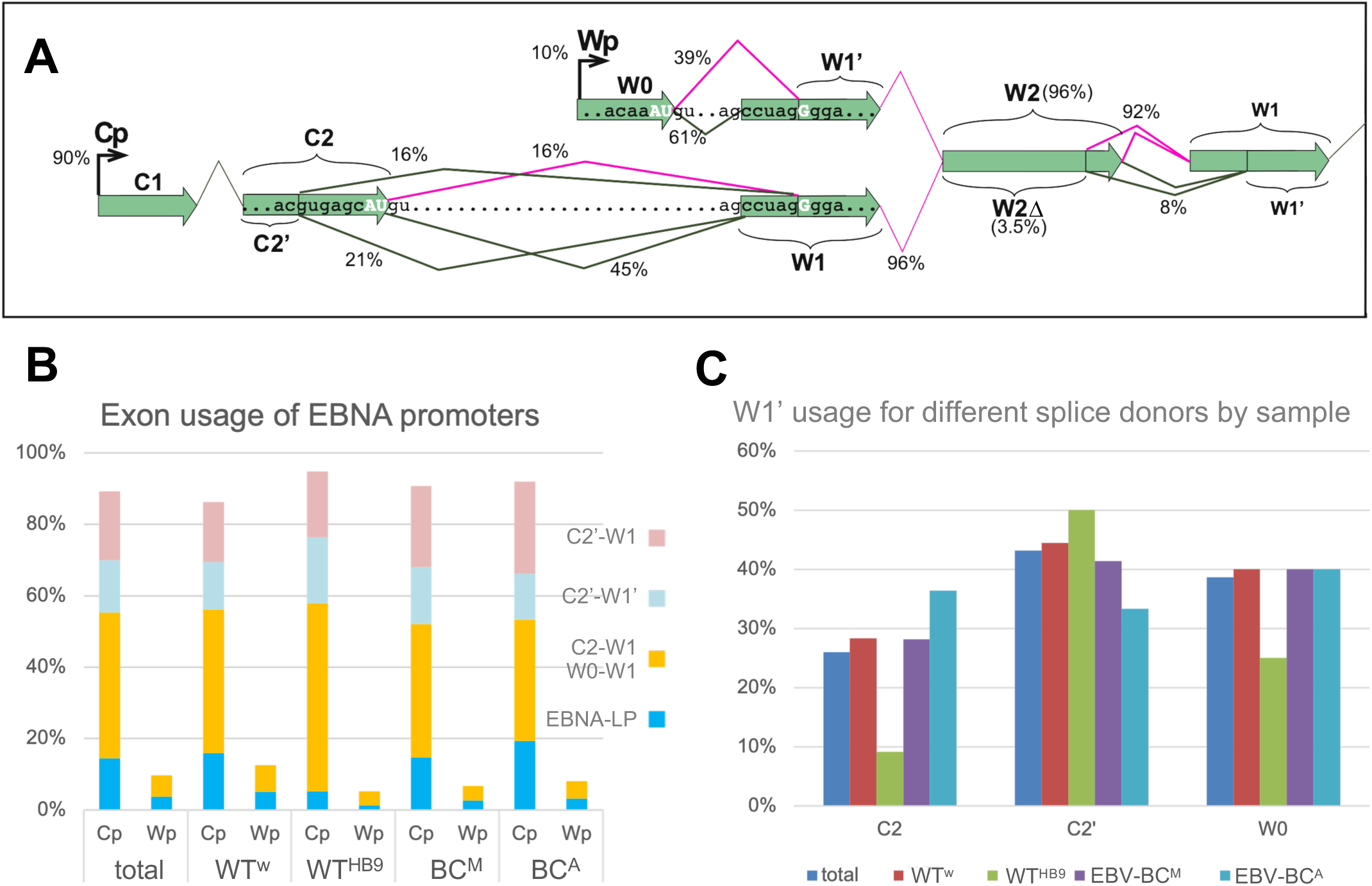
Promoter usage and alternative splicing of the EBNA transcripts. **(A)** Schematic diagram showing the main splice variants and their frequency (expressed as percentage of transcripts from all four samples combined splicing into or within IR1). Magenta splice junctions would retain the EBNA-LP open reading frame **(B)** Graph showing distribution of promoter splice junctions for all combined samples (as A) and separated by the four LCLs analysed. **(C)** The frequency of W1’ usage as the splice acceptor for different splice donors, for all reads combined (total) or individual LCLs made with the different viruses indicated.

Overall, 18% of EBNA transcripts can initiate EBNA-LP translation, although Wp transcripts are more than twice as likely to encode EBNA-LP than Cp transcripts. It is, however, notable that the WT^HB9^ LCL has a much lower frequency – under 7% – of EBNA-LP transcripts (**Figure 2C**). This does not reflect reduced usage of the W1’ splice acceptor, as C2’-W1’ splicing is similar to (or higher than) the other LCLs. Reviewing the reads aligned to the W exons (**Supporting Figure S2**) confirmed the presence of a STOP codon [29] in the first W1 exon of WT^HB9^. This mutation in the first W1 exon from B95-8 was also found in two cDNA clones [31] and in PacBio reads from a B95-8 LCL (data from [30], which observed a similar 8% of Cp transcripts from B95-8 encoding EBNA-LP). This AUG-dependent reduction in transcript abundance, matched by reduced EBNA-LP protein levels in WT^HB9^ LCLs (**Figure 3A** and [29]) suggests translation-associated nonsense-mediated decay of WT^HB9^ transcripts with premature STOP codons.

**Figure 3.**
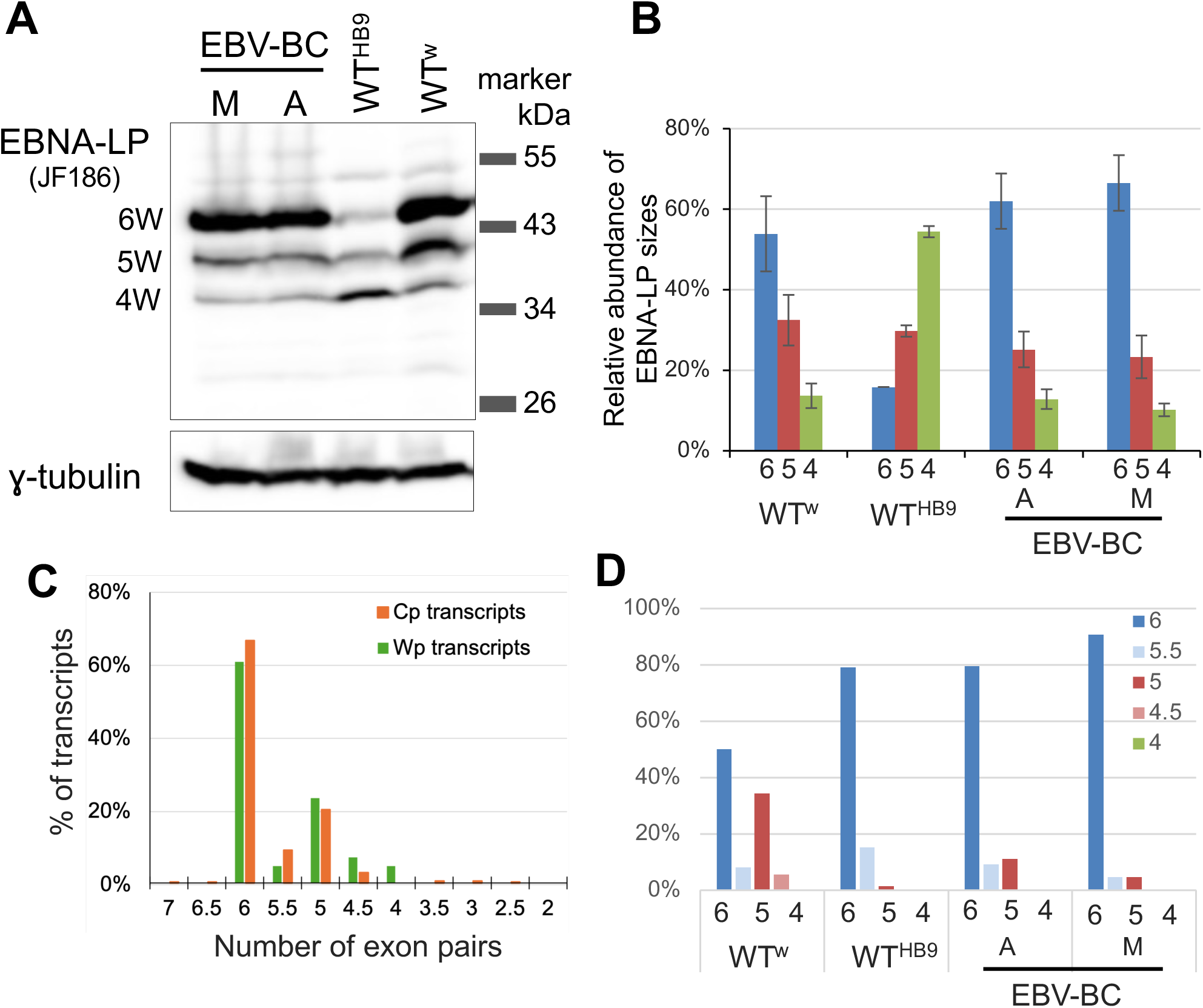
Transcripts spanning IR1 show relatively efficient splicing, and diverse stochastic splicing decisions. **(A)** Western blot of EBNA-LP from the LCLs used in this study. Position and size (kDa) of the marker are shown right. Numerical labels to the left indicate the likely number of W1W2 exon pairs likely in each band size. **(B)** Densitometry of western blot, corrected for the different numbers of antibody epitopes in different length EBNA-LPs, showing relative abundance of EBNA-LP-W4, -W5, and -W6 for each cell line. Number indicates number of W1W2 exon pairs. **(C)** Graph shows percentage of Cp or Wp transcripts with different numbers of W1 and W1’ exons based on counting exons to which nucleotides are mapped, and correcting for total number of aligned nucleotides. **(D)** Distribution of IR1 lengths in Cp transcripts in different cell lines.

### Splicing across IR1 is efficient, and produces a novel EBNA-LP splice variant

To define the origin of the ladder of different sized EBNA-LP proteins commonly seen on western blots, as seen for the four LCLs analysed (**Figure 3A**), we compared protein sizes with transcript structure. The WT^w^ and EBV-BC LCLs produce EBNA-LP protein containing predominantly 6 W repeat domains (EBNA-LP-6W), with progressively less EBNA-LP with fewer repeats, whereas the WT^HB9^ LCL contains very little EBNA-LP-6W, but more EBNA-LP-4W (**Figure 3A**). There are also traces of larger EBNA-LP protein isoforms. Densitometry suggests an approximately 3:1 ratio of 6W:5W in the two EBV-BC cell lines, and a lower 3:2 ratio for WT^w^ (**Figure 3B**). All three lines show an approximately 2:1 ratio of 5 repeat to 4 repeat EBNA-LPs.

We counted the numbers of W1 and W2 exons in reads derived from IR1 to examine how this size variation arises, using the length of reads (in nucleotides) to test the integrity of the alignment of reads to IR1. Around 25% of reads were >85 nucleotides longer than was expected from the number of exons aligned to by the read. This was either because of retained introns (presumably incompletely spliced pre-mRNAs) or a large insertion within a W2 exon, almost always approximately a multiple of the 198 nucleotides of a W1-W2 exon pair (**Supplementary figure S3**), suggesting misalignment of reads to IR1. These misaligned insertions were always found in exon W2, most frequently in the penultimate W2 exon (**Supplementary figure S4A**).

Further analysis also predicted a novel splice donor within exon W2 that lies 21 nt upstream of the canonical splice donor (see examples in **Supplementary figure S3** and schematic **Figure 2A**), producing exon W2Δ – a shorter version of exon W2 – that if in a transcript encoding EBNA-LP, would lack its last 7 amino-acids of its repeat domain. Exon W2Δ was observed in 3-4% of exons aligned to repeats 1-5, and half that (1.7%) for the last repeat exon (**Supplementary table ST3**). In total, 13% of IR1 transcripts contain at least one W2Δ exon. The W2Δ-W1 splice junction was consistently detectable in online short-read RNA-seq data from de novo WT^w^ infection, LCLs and GM12878 RNA-seq datasets, at between 0.7% and 3.1% of the frequency of canonical W2-W1 splice junctions. A similar frequency was observed for W2-W1’ splice junctions, which would frame-shift the EBNA-LP ORF, so it remains unclear whether these alternative splicing events are biologically significant.

To accurately reflect the IR copy number for each read we calculated the sum of nucleotides aligned to IR1 exons (thereby omitting retained introns), relative to the expected nucleotide size of those exons, and multiplied this value by the number of W1+W2 exons we observed to give a final estimate of repeat exon number (**Supplementary table ST3**). These estimates are distributed close to whole numbers of exons in IR1 (**Supplementary figure S3B**), suggesting this approach is a reasonable estimate of IR1 exon usage.

By this measure, the vast majority of Cp transcripts had 5 or more pairs of repeats, while Wp transcripts had 4 or more (**Figure 3C**), and when split by cell line, most of the reads with fewer than 6 W exon pairs are from the WT^w^ cell line (**Figure 3D** and **Supplementary figure S4B**). Rescuing EBV BACs from the LCLs into bacteria, and analysing their restriction pattern, showed that the WT^w^ LCLs had a higher proportion of EBV genomes with a 5 repeat IR1 than the other LCLs (**Supplementary table ST2**), albeit not enough to fully explain the increased 5 repeat transcripts in the WT^w^ LCL. One read had 7 W1-W2 exon pairs: more than expected from the genome sequence, but implied by the larger EBNA-LP isoforms detected on western blots (**Figure 3A** and [29]). Nevertheless together these data imply that number of repeats in transcripts largely matches the number in the underlying genomes, suggesting that exon skipping within IR1 is rare.

### Alignment to IR1 is slightly improved by barcoding first and last repeat

We hypothesised that barcoding the first and last exons, and mapping to the barcoded sequence would improve the fidelity of mapping reads to IR1 exons, by better securing the alignment of the first and last repeat unit. Alignment to the non-barcoded WT^w^ sequence increased the reads skipping exons in the first repeat unit by about 50%, with a corresponding increase in reads aligning to the second repeat unit as their first W exons, and also a slight increase in reads aligning to repeat 5 but not the last repeat (**Supplementary Table ST4**). However, barcoding did not reduce the overall number of mis-mapped large insertions, although it did alter their positioning (**Supplementary Table ST5**). Thus the fidelity of alignment to homogeneous repeats is only marginally improved by barcodes in first and last repeats.

### Defining full-length transcripts reveals RNA processing sites

In principle, transcript abundance can be estimated from nanopore dRNA-seq data by either measuring frequencies based on either polyadenylated 3’ ends; from defined full length reads; or from any reads that span the region being analysed. We initially extracted ‘full-length’ reads based on the expectation that authentic 5’ and 3’ ends would be present in at least 5 transcripts. This identified the main viral latency promoters (Cp, Wp in repeats 1 and 2, LMP1, LMP2A and BARTs) and polyA sites, but also a number of non-canonical 5’ ends (**Supplementary table ST6**). Indeed, over 96% of reads contained a reproducible or previously annotated polyadenylation site [32], whereas under 60% of reads had a biologically explicable 5’ end (**Supplementary table ST7**), such as those previously defined by CAGE-seq [33].

Non-canonical 5’ ends were particularly frequent across the GC-rich repeat sequences of EBNA1 and EBNA2, and the 3’ UTR of LMP1, suggesting these regions are more frequently break, or have RNA structures that can block the nanopore, curtailing sequencing. Conversely, a large number of reads with a 3’ end at the annotated BHRF1 polyadenylation site, had a 5’ end close to the end of miR-BHRF1-2 (and much more rarely miR-BHRF1-3 or miR-BHRF1-1) (**Supplementary table ST7**). The much higher abundance of ends generated by miR-BHRF1-2 cleavage may have implications for the order in which BHRF1 miRNAs are processed, while reads polyadenylated at BHRF1 and cleaved at the site of mirBHRF1-1 processing suggest that at least some BHRF1 miRNA processing precedes RNA splicing, rather than miR-BHRF1-1 excision from the intron (**Figure 4A**).

**Figure 4.**
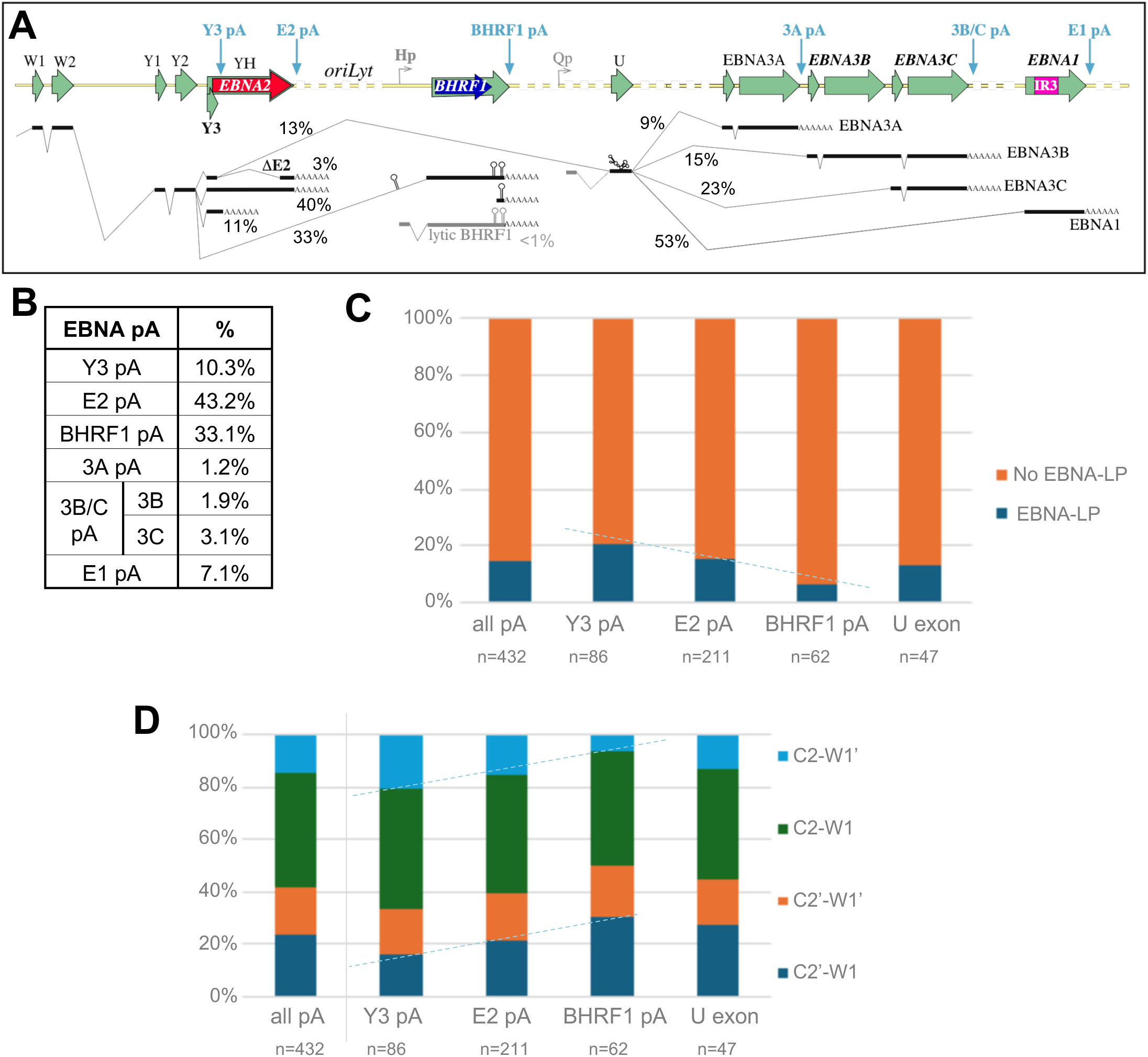
Frequencies of EBNA transcript polyadenylation and alternative splicing events. **(A)** Schematic representation of the main Cp/Wp-derived RNAs observed or inferred from the nanopore reads in this study. Blue arrows indicate poly adenylation sites. Reads/exons in grey are previously observed splicing events that are infrequent or absent in these data. **(B)** Table of relative frequencies of transcript pA sites inferred to arise from Cp or Wp promoters (EBNA transcripts). EBNA3B and EBNA3C relative abundance within the EBNA3C pA group was inferred from the ratio of reads extending upstream of 3C Exon1. **(C & D)** Graphs testing co-ordination between alternative splicing of promoter exons and the polyadenylation site usage, shown either **(C)** in terms of whether the splicing generates the AUG start codon for EBNA-LP, or **(D)** frequency of different Cp splice combinations for each pA site. All pA sites downstream of the U exon have been merged into one group. Dashed lines indicate trends noted in the article text.

Characterising the 3’ ends of transcripts confirmed the known polyadenylation sites of EBNA, LMP and BART transcripts [32], and identified a novel EBNA transcript polyA site between the Y3 splice donor and the EBNA2 start codon (herein called Y3 pA) (**Figure 4A**). EBNA transcripts most commonly used the canonical EBNA2 or BHRF1 polyA sites, with Y3 pA used in 10% of transcripts (**Figure 4B**). Since a third of EBNA transcripts use the BHRF1 pA site (and lytic BHRF1 transcripts were rare), we can infer that – at least in LCLs – BHRF1 transcripts are a fundamental part of the latency gene expression profile of LCLs, despite the BHRF1 protein being barely detectable. EBNA3C pA reads whose 5’ end was upstream of the EBNA3C exon 1 splice acceptor (n=26) showed 5:8 ratio (approximately 60:40 split) of 3C:3B-encoding transcripts. Combining this with pA site usage (assuming all RNAs are equally stable) allows inference of the ratio of different splicing events downstream of the U exon, suggesting that more 3’ EBNA gene transcripts are progressively more frequent (**Figure 4A & 4B**), suggesting the U exon splice donor more frequently uses the more distant acceptor, though whether this is determined by splicing or polyadenylation mechanisms cannot be distinguished. Interestingly, the pA tails at BART and BHRF1 pA sites (the transcripts associated with miRNA processing) were around double the length of other pA tails (Supplementary table ST6).

We did not observe any of these downstream EBNA transcripts initiating at the latency II/I-specific Qp promoter, but cannot formally exclude its usage due to the very high breakage rates of EBNA1 and EBNA3 transcripts (**Supplementary table ST7**). Use of latency polyadenylation sites outnumbered lytic gene pA sites by 9:1, with 2/3 of the lytic cycle pA sites being from BHLF1 transcripts (**Supplementary table ST6**), suggesting reactivation is occurring in a small proportion of cells, but we do not explore these further as they have been studied in detail elsewhere [2, 34].

### EBNA promoter, polyadenylation and splice site usages appear to influence each other

The alternative splicing of the EBNA transcripts has a wide range of options, and we aimed to test the hypothesis that there is co-ordination between promoter usage, polyadenylation, and alternative splicing. Certain splice acceptor-donor combinations appear to be preferred, with Y2-BHRF1 being hugely preferred to Y3-BHRF1, and conversely the U exon acceptor was spliced exclusively to Exon Y3 in our data, even though Y2-U splice junctions have previously been observed elsewhere [30, 35, 36].

We next assessed whether different polyadenylation signals are associated with different proportions of EBNA-LP initiating transcripts. Wp-initiated transcripts were consistent (albeit with low read numbers), with an approximately 50:50 use of W1 and W1’ regardless of pA site. Shorter Cp transcripts appeared generally more likely to encode the EBNA-LP start codon than longer ones, although this trend slightly reversed in U exon transcripts (**Figure 4C**). This change is associated with progressively increasing use of C2’/W1 splicing and reduced C2/W1’ in the EBNA2-pAand BHRF1-pA transcripts relative to those with Y3 pA. The other two splice junctions are unchanged in frequency (**Figure 4D**). We tested whether increasing distance of the pA site from Cp is statistically associated with either increasing use of C2’ over C2 (p=0.078), or decreasing propensity to encode EBNA-LP (p=0.0521) using a Cochran–Armitage test for trend. While this does not quite reach conventional boundary for statistical significance, it does suggest that there is likely to be co-ordination between splice site usage and pA sites that would bear further study.

### The BART transcripts in the EBV BAC interact with inserted expression cassettes

The deletion in B95-8 removes BART Exons II to IV (including the RPMS1 start codon), and retains only four BART miRNAs from cluster 1 (**Figure 5A**) and miR-BART-2 (between exon IV and V). We therefore assessed how this deletion and the inserted BAC expression cassette (lying between the BARTs’ promoter and downstream exons) affected transcripts from this region. Antisense to the BARTs, nearly 400 reads used the GFP polyadenylation site were detected, compared to just 11 for the SV40 pA used by the hygromycin phosphotransferase cassette (**Supplementary tables ST7)**. Despite the large number of GFP reads, when viewed by fluorescence microscopy, only a small proportion of the cells are visibly green, which suggests that many of the GFP reads may derive from a small percentage of the cells. The BART transcripts predominantly used either the canonical site at the end of BART exon VII [20], which confirms that BARTs can still be transcribed across the BAC insert, or the SV40-derived pA sequence between the Hygromycin and GFP cassettes (**Supplementary table ST7** and **Figure 5A**). In total over 200 reads use one of these pA sites (that we collectively call BART transcripts), of which one third reach Exon VII, while two thirds terminate in the SV40-derived pA used to terminate GFP transcripts.

**Figure 5.**
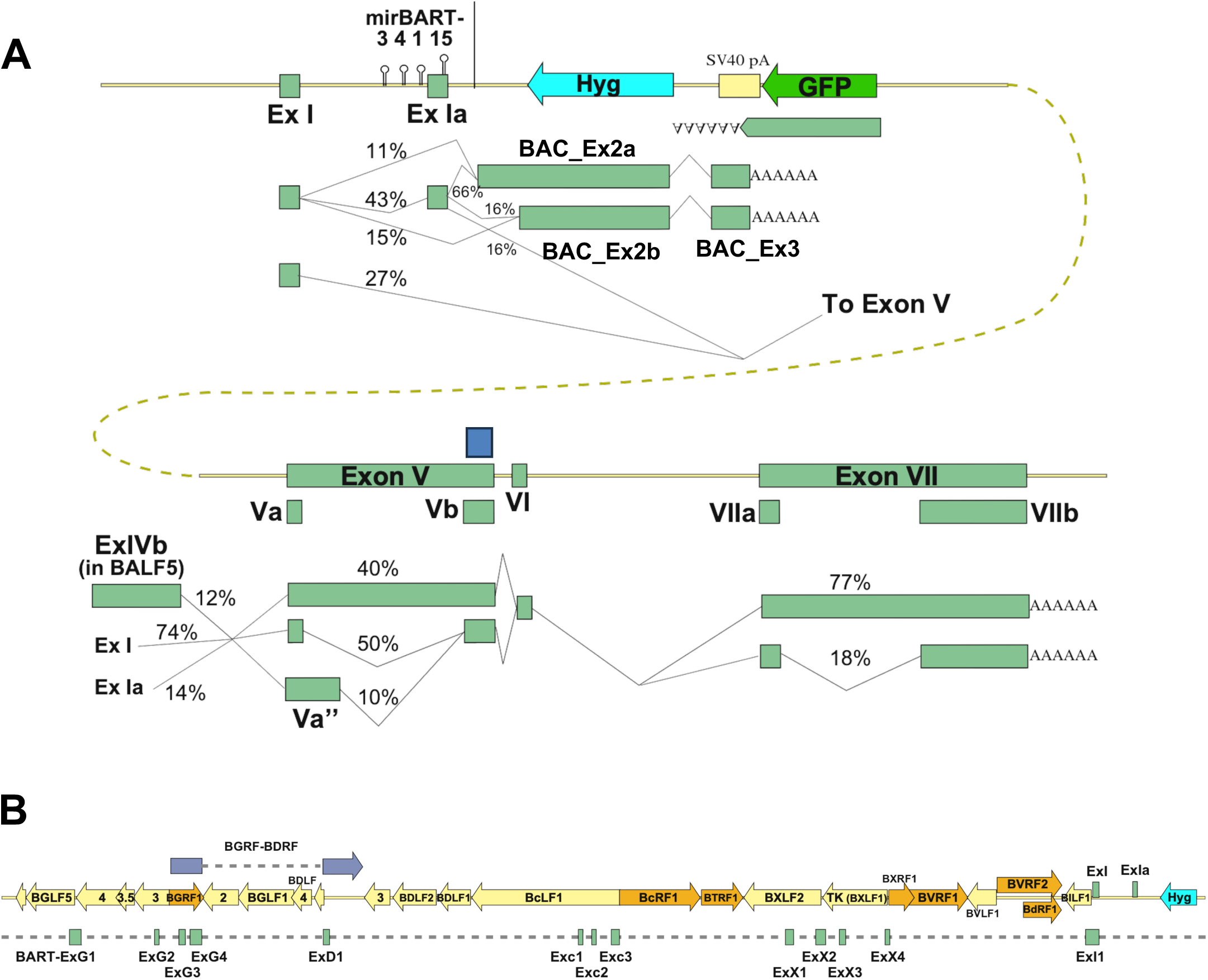
The splicing profile of BART transcripts of the EBV BAC. **(A)** Schematic representation of transcripts from the BART promoter spanning the EBV BAC. Percentage values are in vertical groups, and indicate the proportion of differential splice donor, splice acceptor or exon usage events. The Hyg transcript is omitted because only one read supported its existence. **(B)** Novel exons identified upstream of the canonical BART promoter (ExI) are shown as rectangles below the EBV genetic map. Novel exon names are based on the ORFs in which they are found. ORFs are colour coded by sense (orange) or antisense (yellow) orientation.

The short BART transcripts using SV40pA start with Exon I and usually (60%) Exon Ia, splicing to either of two cryptic splice sites, within either the truncated mirBART5 sequence or 60 nt before the end of the antisense Hygromycin phosphotransferase (Hyg) ORF (**Figure 5A**). Both of these exons (BAC_Ex2a and b respectively) extend across all but 20 nt of the Hyg ORF, then splicing to another exon (BAC_Ex3) just past the SV40 promoter. The GFP and BAC_Ex3 polyadenylation sites are those used by the bidirectional transcripts of the SV40 polyomavirus, having 3’UTRs with 80 nt of reverse complementarity to each other. **(Figure 5A).**

The longer BART transcripts – ending in the canonical exon VII – exhibit alternative splicing in exons V and VII that have been previously reported [37]. We saw usage of exons V, Va and a 281 nucleotide exon (Va’’) [38], but no examples (in 41 reads) of exon V’ required to encode BARF0 [37], which implies (p<0.05) that exon Va’ has no more than 7% abundance in LCLs. Similarly there was no evidence for transcripts initiating close enough upstream of hypothesised ORFs A73 or RPMS1 for their translation.

### EBV transcripts can extend between distinct transcription units, spanning the EBV genome

As well as these canonical BARTs, ≈15% of BART reads extend 5’ of the canonical BART promoter (**Supplementary table ST8**). These reads contain a collection of exons from 20 kb upstream of the BARTs, that connect to the BARTs via a novel splice acceptor around 25 nt upstream of the BART Exon I transcription start site. Many of these are antisense to established late gene ORFs, but some arise within the BGRF-BDRF transcript region (**Figure 5B**). The novel exons could encode possible alternative versions of the BGRF1-BDRF1 terminase large subunit, and three other small ORFs, but whether these are translated will depend on the promoter usage of these transcripts, which may differ depending on cell type and latency transcription programs. In these LCL transcripts, there is no new promoter, but rather the few with long enough sequences are spliced from Cp, W repeat or U exons within EBNA transcripts.

For the LMP2 transcripts, there is a 3:1 ratio of the LMP2A:2B splice junctions, but 2.5% of splice donors to LMP2 Ex2 come from exons further upstream (**Supplementary table ST8**): half are from a novel splice donor within BART Exon VII, but others use BART Exon I, BAC_Ex2, and EBNA U exon splice donors. Overall, around 1% of reads with a Cp or Wp initiation site terminates in a canonical polyA site downstream of EBNA1. Because these longer reads are more likely to fragment, it is unclear how common they truly are, but does suggest that either RNA polymerase can extend transcripts for the full length of the EBV genome from Cp to the LMP2 polyA, or that compound EBV reads can arise from trans-splicing at a detectable level. Whether such reads are biological noise, or hold relevance to EBV biology remains an open question.

## Discussion

The transcriptome of EBV has been studied for over 40 years, from cosmid cloning and sequencing of the B95-8 EBV genome [39], followed by Northern blotting, S1 mapping, sequencing of cDNA clones [39, 40], and subsequently updated to the composite EBV annotation NC_007605 that is the current EBV reference [41].

Additional details have been revealed by RNA-seq of latency III B95-8 [42] and reactivated Akata and Mutu EBVs [34], and mapping of lytic cycle transcription start [43] and polyadenylation sites [32], and recent long-read sequencing of lytic cycle cDNAs [2, 44], to mapping of viral circular RNAs [45].

### Novel insights into the EBV latency transcriptome

This long-read direct RNA transcriptome of EBV latency transcripts defines a new latency polyA site near exon Y3, short version of Exon W2, the altered BART splicing of the B95-8 BAC, and previously unreported exons mapping in between EBNA1 and BARTs, as examples of long transcripts that can span conventional gene boundaries. Similar transcripts extending from the BARTs into LMP2 [2] were identified in the EBV lytic cycle, although those transcripts were characterised by extensive exon skipping in LMP2, in contrast with the canonical LMP2 splicing in our extended transcripts. Intriguingly, long transcripts spanning much of the virus genome were also seen in a reactivated KSHV transcriptome [46] although these were generally 2 or 3 exon transcripts. While some of these long distance splice junctions were detectable during KSHV latency, it is unclear whether complex EBV-like transcripts are also made by KSHV during latency, and whether they are biologically important. Since our (and other long-read herpesvirus transcriptomes) analysed total RNA, is unclear which RNAs are exported to the cytoplasm for translation, are functional nuclear lncRNAs, or are incompletely processed or aberrant RNAs with no coherent function. Mechanistically such transcripts could arise through either read-through transcription (inefficient polyadenylation), as is seen is “Downstream-of-Gene” ncRNAs [47], or trans-splicing, the post-transcriptional joining of distinct pre-mRNAs [48].

The complexity of the EBNA transcripts can only be addressed with long read approaches. The vast majority of Cp transcripts include all IR1-derived exons, suggesting exon skipping across IR1 is rare, although there remains uncertainty because the IR1 repeat number is not entirely stable. Conversely, Wp-initiated transcripts, although they often initiate at the Wp promoters closest to Cp in these LCLs, vary more in length than Cp transcripts. This implies that the majority of EBNA-LP deriving from Cp will correspond to the maximum size for that genome, while Wp activity likely contribute the smaller EBNA-LP isoforms. Additionally, since Wp more often generates EBNA-LP transcripts than Cp, Wp transcripts likely produce a disproportionate amount of the cells’ EBNA-LP. Since Wp is the first promoter activated after infection [49], its higher production of EBNA-LP suggests that EBNA-LP protein production is more important in the early stages of infection, fitting both EBNA-LP’s early detection after B cell infection (alongside EBNA2) [50, 51], and its importance for transcription of other viral genes [3], likely by preventing Sp100 and Sp140L from repressing un-transcribed regions of the EBV genome [4].

Similarly, BHRF1 transcripts are made early after infection, but after the first few days post infection, BHRF1 protein becomes very hard to detect by western blotting [52]. Our analysis does not provide an easy explanation for why this might be: BHRF1 transcripts from Cp are less likely to contain an EBNA-LP AUG than those terminating at Y3 or EBNA2 polyA sites (**Figure 4D**), so competition with EBNA-LP translation is unlikely. Perhaps skipping exon Y3 makes the cryptic AUG from exon Y2 more accessible (by changing the mRNA secondary structure), thereby suppressing downstream translation of BHRF1. Given the abundant fully processed BHRF1 transcripts, it seems unlikely that miRNA processing (representing only 15% of transcripts, albeit probably more short-lived) would be sufficient to so substantially reduce translation. Together this suggests that the suppression of BHRF1 protein synthesis in latency III cells may lie downstream of RNA processing. Nevertheless, miRNA processing was evident in 15% of BHRF1 reads, most of which were truncated at the 3’ cleavage site mirBHRF1-2. Since expression constructs with a mutated mirBHRF1-2 hairpin exhibit loss of mirBHRF1-3 production [53], this suggests that mirBHRF1-2 processing is a pre-requisite for (and precedes) mirBHRF1-3 processing, and that after mirBHRF1-3 processing the RNA is rapidly degraded.

BHRF1 transcripts containing W1-W1 splice junctions reportedly produced during the lytic cycle of Akata and Mutu EBV [2] were absent from our latency III transcriptome. Instead, >99% of BHRF1 transcripts spliced from exon Y2 to BHRF1 exon 2, consistent with previous analyses [30, 52]. This contrasts with splicing to the U exon, where ≈90% of transcripts also include exon Y3 (similar to the 84% seen by PacBio sequencing [30]). Thus both Y2 and Y3 splice junctions are key in defining alternative splicing of EBNA transcripts.

We have also characterised the disruption of BART splicing that results from the introduction of a BAC into the B95-8 genome. Both the Hygromycin phosphotransferase and GFP expression cassettes use pA sites from SV40 polyomavirus. The pA downstream of the Hygromycin resistance gene contains only the SV40 T antigen polyA site, whereas the longer region of SV40 after GFP also contains the antisense pA used by the SV40 late transcripts. It is this antisense pA that terminates 2/3 of the BART transcripts prematurely. It is perhaps surprising, however, that the cryptic exons antisense to Hygromycin are never incorporated in longer BART transcripts. This suggests that there are evolved RNA structures or RNA-binding proteins that support the correct processing BART transcripts using the canonical exon VII pA.

### Improving EBV genome annotation

One of the goals of this project was to create an EBV transcriptome annotation that captures the most common latency transcript isoforms (downloadable for the B95-8 BAC from ebv.org.uk). The relatively efficient IR1 splicing of Cp transcripts supports a simplification that Cp transcripts contain all IR1 exons. However, the variable Wp usage and new W2Δ exon (being distributed throughout IR1) are challenging to annotate without creating an unreasonable number of distinct mRNAs. The identification of transcripts spanning almost the entire genome led us to use oriP as the “ends” of the EBV genome, rather than the terminal repeats used in NCBI annotations, which breaks LMP2 transcripts. Although LCLs had no reads in oriP, these were observed during lytic reactivation [54]. Thus, in the absence of RNA-seq aligners able to map to circular genomes, future studies will need to tailor annotation files and analysis pipelines to specifically capture the biology they are investigating. To support improved transcriptome annotation of EBV for RNA-seq, we have produced updated GFF3 file for botht eh B95-8-BAC and the NC_007605_M (the prototype EBV genome shifted to start at the MluI site in oriP) that assumes EBNA transcripts have 6 repeats initiating at Cp, and including the BHRF1 and Y3 pA sites, and also a new ‘gene’ annotation for the spliced region upstream of BARTs. Exons C2’, W0 and W2Δexons are annotated as orphan features for reference (Supporting data file SD1).

### Limitations, future optimisation and utility of dRNA-seq of EBV

Despite the advantages of direct RNA sequencing, which avoids biases-introduced through PCR, and captures long-range associations between promoter usage, polyadenylation, and splicing, we identified shortcomings and biases even in this minimally manipulated direct RNA sequencing data.

We wrote a script to compensate for the failure of minimap2 to correctly align the short 5’ W0 exon, while imprecise splice site assignments (also reported elsewhere [55]) required us to use reference-guided alignment, but could not resolve the position of the W2Δ splice donor. While these will be improved by better base-calling accuracy, the miscounting of W exons due to incorrect alignment to the IR1 repeat require strategies like the nucleotide counting used herein to resolve, until aligners’ handling of repetitive sequences improves.

Other inaccuracies stem from sequencing being terminated before the canonical 5’ end of the transcript. This was particularly prevalent in EBNA1, but also in GC-rich regions of EBNA3B/C, EBNA2 and BHRF1 transcripts. The Gly-Ala repeat in EBNA1 forms G quadruplex structures [56], suggesting that RNA structures blocking pores (rather than hydrolysis of the RNA’s phosphate backbone) is responsible for regions of increased sequencing termination. This could be reduced by using a reverse transcriptase with better processivity, such as Induro (NEB), which improves the proportion of intact 5’ ends of transcripts [57], and is now recommended by Oxford Nanopore Technologies. This, alongside improved flow cells and base-calling, should produce even more robust dRNA-seq data going forward.

The complexity and low abundance of EBV latency transcripts makes it particularly challenging to fully characterise. Alongside improved RNA integrity, the small proportion of cellular reads originating from EBV limits the study of EBV transcript complexity. To increase the number of EBV-derived reads requires development of enrichment strategies such as targeted library preparation, hybrid capture strategies analogous those used for EBV genome sequencing [58] but suitable for maintaining intact RNA transcripts, or adaptive sampling during sequencing [59].

The polyA-based sequencing means that EBV’s many abundant but non-coding RNAs are not detected in our data-set. These include 1.3kb and 0.9kb RNAs from BHRF1 liberated from BHRF1 transcripts by miRNA processing, that are abundant during the EBV lytic cycle [15, 53]; the RNA-pol III-transcribed EBERs; the putative stable intronic sequence RNAs (sisRNA-1 and -2) that arise from the introns between repeating W1 and W2 exons [14]; intronic RNAs spanning the terminal repeats that work with EBER2 to regulate LMP2 [60]. Detecting such RNAs in direct RNA-seq would require specifically targeted adapters, as used for direct rRNA sequencing (e.g. [61]).

Nevertheless, direct RNA whole transcript sequencing promises to provide the most comprehensive and least biased snapshot of the EBV transcriptome. It will be important to determine differences between lab strains and authentic circulating EBVs, and validate whether genetic modifications have unexpected impacts on transcript processing, as seen for an accidental splice acceptor, and elevated EBNA2/LP and LMP1 protein expression that characterised two mirBHRF1-mutant viruses [30], and the BAC-induced alterations in BART splicing and polyadenylation. Similarly in this study we detected a reduced prevalence of the C2-W1’ splice junctions in WT^HB9^ that is also observed in B95-8 LCLs [30]. Since this splice junction forms the EBNA-LP start, but B95-8 and WT^HB9^ EBVs encode a STOP codon in what we now know is the first W1 exon, we suggest that the reduced prevalence of C2-W1’ splice junctions is a signature of nonsense mediated decay of transcripts due to the initiation and premature termination of EBNA-LP translation..

Interpreting the transcriptome of EBV is further complicated by the recent discovery that LCLs are not homogeneous, but rather represent a collection of cells with diverse B cell differentiation states [62, 63]. It remains possible that some EBV transcript variants are tied to particular B cell differentiation states (influencing, or influenced by specific aspects of B cell biology), rather than simply being the stochastic variations in splice site usage. Truly understanding the nature and control of EBV transcription will require a combination of sorted or single cell techniques with targeted long read sequencing.

## Materials and methods

### Generation of recombinant EBV containing modified W2 exons

To generate EBV with a ‘barcoded’ IR1, the Mfe I/Pci I restriction fragment from IR1, which contains the W exons, was cloned into a pBR322-based plasmid. The plasmid was amplified by PCR (primers in **Supplementary table ST1**) and circularised by InFusion cloning (Takara Bio), introducing pairs of synonymous SNPs (creating either Mlu I or Pvu I restriction sites) in exon W2. The Mfe I/Pci I restriction fragments were cloned into either p8359 (Mlu I barcode) or p8342 (Pvu I barcode), which were combined by Gibson Assembly (NEBuilder HiFi Assembly mix) with wild-type BamW fragments to assemble an IR1 repeat array as described previously [3], with the Pvu I SNPs in the first repeat unit, and the Mlu I SNPs in the last repeat unit. This IR1 repeat array was recombineered into the B95-8 BAC (WKO.4 from [3]) using RecA-based recombineering [64]. The resulting EBV-BAC with a barcoded IR1 (pHB9-W4-MR11+4.27 – herein called EBV-BC) was purified and transformed into HEK293 cells to generate EBV-producing cell clones (-AJ2 and -MJ2) as previously described [3].

### Cells and viruses

Virus production was induced from EBV-producing 293 cell clones carrying episomal EBV genomes as described previously [3]. Specifically, virus was isolated from the two barcoded B95-8-BAC producers (293-MR11-AJ2 and -MJ2); from 293-HB9-A4 containing our clone (WT^HB9^) of the p2089 B95-8 BAC [1], whose IR1 contains 6.6 repeat units [29]; and 293-WT^w^.1-A11 [3], which is WT^HB9^ engineered with a homogeneous IR1 repeat to produce WT^w^ [3]. These viruses were used to infect B cells isolated (RossetteSep B cell purification kit) and pooled from buffy coat residues of two anonymous blood donors, and grown into LCLs in RPMI supplemented with 10% FCS and 4mM L-glutamine.

### RNA extraction and sequencing

LCLs were seeded at 3×10^5^ cells/ml 24 hours prior to harvesting cells. RNA was extracted using TRIzol, followed by two additional washes in 70% ethanol prior to resuspension, and polyA-based library preparation with RNA002 kit (Oxford Nanopore Technologies) including reverse transcription to stabilise the RNA (using Superscript III; Life Technologies), followed by direct RNA sequencing using MIN106D version R9.4.1 flow cells. FAST5 files were base-called to generate FASTQ files using Guppy (version 6.1.5) [65].

### Transcript mapping and characterisation

Specific commands used for the processing and mapping of FastQ reads are available as a script (see Data Availability below). To summarise, reads were aligned separately for each cell line in an annotation-aware manner, to either the corresponding exact EBV-BAC sequence (“vs self”) or to WT^w^ EBV-BAC sequence (all identical IR1 repeat units) with minimap2 (v2.24) [66, 67]. Where not specified, analyses were performed on the latter “vs self” alignment. Manipulations of SAM and BAM files were performed using SAMtools (v1.3.1). For pooled analyses, SAM files were concatenated. IDs of EBV-aligned reads were extracted from the SAM file and used to extract EBV-specific FAST5 data (fast5_subset command within ont-fast5-api v4.1.1). An in-house script (W0_correcting.r) identified W0 exon-derived sequences that had been removed by soft-clipping during the initial alignment by Minimap2, and modified the SAM files to appropriately align these W0-derived sequences. Common polyadenylation (pA) sites and transcription start sites (TSS) were identified and reads sorted into full-length and various incomplete/rare read groups using an in-house script (transcript_classifying.r). Full length transcripts are defined as those whose 5’ and 3’ ends are both no more than 20 nucleotides away from a reproducible start or stop position. Reproducible start and stop sites are where a 5’ or 3’ end occurs in at least 5 reads within a 5 nt window. Next a third in-house script (IR1_repeat_counting.r) was used to extract the number of IR1 repeat units to which reads were aligned, and the number of nucleotides aligned to each IR1 W exon in each repeat unit (including the novel W2Δ exon identified herein). Information about polyA lengths was extracted using nanopolish (v0.14.0) [68]. Reads were then parsed using previously published perl scripts (classify_transcripts_and_polya_segmented_V2.pl and all_transcripts_grouped_by_polya.pl [69]) to extract and collate 5’ end, 3’ end, and splice site co-ordinates from each read, to filter reads according to their pA tail status, and adjust the 5’ end position to correct for the ≈10 most 5’ nucleotides, that are lost by nanopore sequencing [69, 70]. Subsequent characterisation and quantitation of splice junctions were conducted using excel formulae to extract details from these outputs. IR1 analyses were conducted on full-length reads only. Other transcript analyses used the polyadenylated subset of reads to define splice donors and acceptors, but used all reads (including those without detectable polyadenylation) to quantify splice site usage. Read alignments were visualised using Integrated Genome Viewer (IGV) v2.16.2. Statistical analysis to assess whether later polyadenylation was co-ordinated with changes in Cp alternative splicing outcomes used the Cochran–Armitage test for trend (function: `prop.trend.test` in R version 4.5.0). Putative W2-W1 splice junctions were validated in short read RNA-seq data for de novo WT^w^ infection (SRR31410412, SRR31411669, SRR31411671), arbitrary B95-8 LCLs (SRR16011070, 71, 81, 91), and GM12878 cells (SRR23957814) using the Sequence Read Archive filter feature to count reads containing sequences matching the 20 nucleotides centred on the splice junction.

### Western blotting

Immunoblots were performed using RIPA protein extracts and mini-Protean gel electrophoresis and blotting equipment (Bio-Rad) to transfer the gel contents to nitrocellulose as described previously [29] and imaged using an Azure 600 chemiluminescence imager. The anti-EBNA-LP JF186 antibody hybridoma supernatant was used at 1:100; Mouse anti-ɣ-tubulin (clone GTU-88; Sigma-Aldrich) was used at 1:1000.

### Analysis of EBV episomes

To extract EBV genomes from stably infected cell lines (“episome rescue”), DNA was extracted from approximately 3×10^6^ cells using the low molecular weight DNA extraction protocol [71], with additional use of MaXtract tubes (Qiagen) to maintain separation of phases during the phenol extraction step. DNA was resuspended in 50 µl of TE-RNase, of which 2 µl was electroporated into NEB10-beta bacteria using a Bio-Rad Gene Pulser. BAC DNA was extracted chloramphenicol-resistant colonies by alkaline-lysis miniprep, digested with EcoRI and analysed by pulsed field gel electrophoresis (Bio-Rad CHEF-DR II) to visualise the IR1 band (largest fragment).

## Supporting information

Supplementary tables and Gff3 files

## Data availability

Perl scripts from [69] are available with documentation from zenodo.org/record/7101768. In-house scripts with documentation is available from github.com/robertewhite/ebv-transcriptomics-tools. The Bam files for reads aligned to EBV (with modified W0 alignment), the FAST5 and FastQ files for EBV reads are available in the EMBL Nucleotide Archive (Study accession number: PRJEB83447 – ERR14129300-3). All raw and processed EBV data from this study, including the NC_007605_M and EBV-BAC genome annotations, EBV plus cellular FastQ reads, and scripts are linked from ebv.org.uk.

## Acknowledgements

Thanks to Claudio Elgueta for providing the B cells used to produce the LCLs described in this project. Thanks to Micah Luftig for unpublished observations from PacBio sequencing from [30]. Thanks to Paul Farrell for critical reading of this manuscript.

## Funding

REW and WW were supported by MRC grant MR/S0022597/1. AM-L, IZ, MR, JH were supported by Imperial College MSc project stipends. AG was supported by Wellcome Trust (grant WT099273/Z/12/Z) to Prof Martin Allday. DAM and ID-B were supported by BBSRC grant BB/M02542X/1.

## Author contributions

Conceptualisation: REW, AM-L, IZ

Computational Biology and programming: REW, AM-L, IZ, DAM

Data Analysis REW, AM-L, IZ, JC, HIC

Transcriptome annotation: REW, AG, MS

Experimental Work: JH, MR, WW, ID-B

Manuscript preparation: REW, AM-L

## Conflict of interest statement

The authors have no conflict of interest with the contents of this manuscript.

## Supporting data – Figure and table legends

**Supplementary table ST1:** Oligos for barcoding IR1

**Supplementary table ST2:** Episome rescue W repeat totals

**Supporting table ST3: Calculation of IR1 number of repeats.** In-house script IR1_repeat_counting.r was used to count the number of nucleotides aligned to each of the latency exons. Only reads with C1 or C2 exons, plus those previously processed for W0 correction are analysed. Calculated W1-W2 exon number compares the observed number of nucleotides aligned to W exons with the number expected for the count of exons to which nucleotides are aligned (column O), and uses that to calculate a repeat number corrected for alignment problems (column P). Highlighting indicates mismatches between the number of W1 and W2 exons, or between observed and expected repeat number.

**Supporting table ST4: Reads skipping barcoded exons in vs self and vs WT^w^ alignments.**

**Supporting table ST5: Number of oversize W2 exon alignments by sample.**

**Supporting table ST6: 5’ and 3’ positions used to define full length reads.** Unbiased identification of "full length" reads defines promoters and pA usage, but 5’ ends also include sites of frequent read breakage in EBNA2 and EBNA1 genes.

**Supporting table ST7: Read integrity separated by different promoter and polyA site usages.** This table combines all canonical 5’ and 3’ ends identified in table ST6, plus all sites that correspond to previously described promoters or polyA sites. Total usage is the read count for reads with that pA site or 5’ end. % full length represents % of reads by pA site that also contain a defined 5’ end (left two tables); or 5’ ends that also contain a defined pA site (right tables). These data are summarised in the tables below that show how frequently a read has a canonical 3’ (left) or 5’ end, showing that (as expected for the pA-initiated sequencing method) that the vast majority of reads have canonical 3’ ends (pA sites) but under half of sense reads run all the way from pA site to promoter.

**Supporting table 8. Sense orientation splice junctions.** Includes only sites that are frequent in our data or previously annotated. Numbers in cells are number of reads with that junction. Donors marked undefined do not have a consistent exon.

**Supporting Data SD1. GFF3 files for annotation of the B95-8 BAC and prototype EBV genome NC_007605.** Note that the sequence of the

**Supporting Figure S1.**
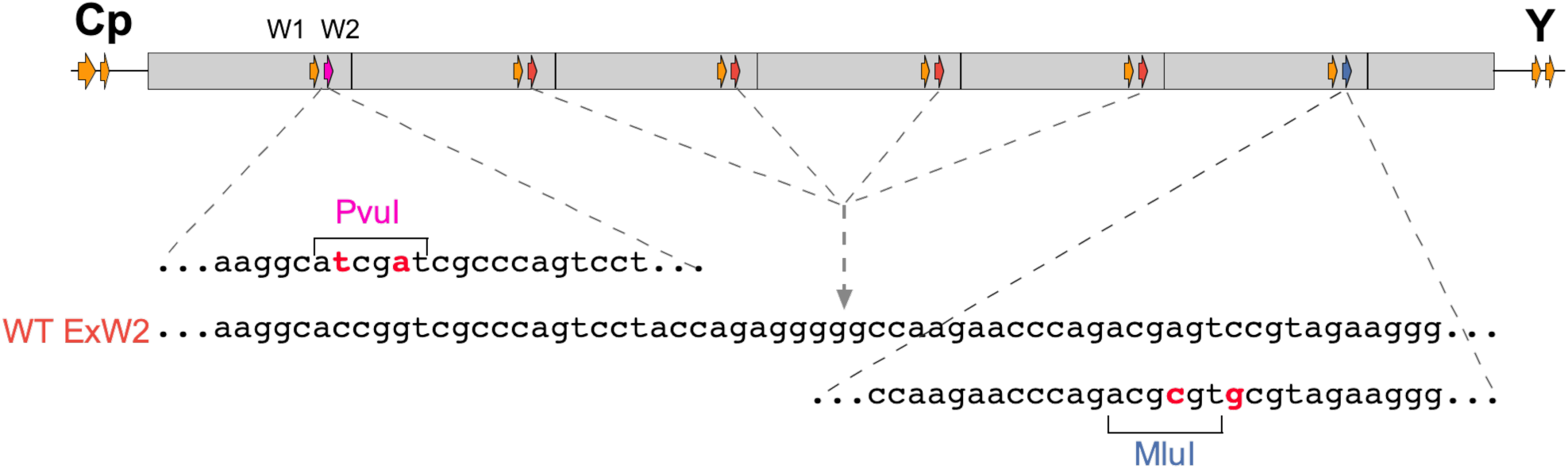
Barcoding sequences of EBV-BC. Schematic representation of the IR1 repeat indicating the Cp, W1 and Y exons (Orange), with the W2 exons coloured according to their sequence, which is shown below: With the first W2 exon PvuI barcoded (Magenta; top sequence); the next four having Wild-type Exon W2 (Brick Red; middle sequence) and final W2 exon MluI barcoded (Blue; bottom sequence).

**Supporting Figure S2.**
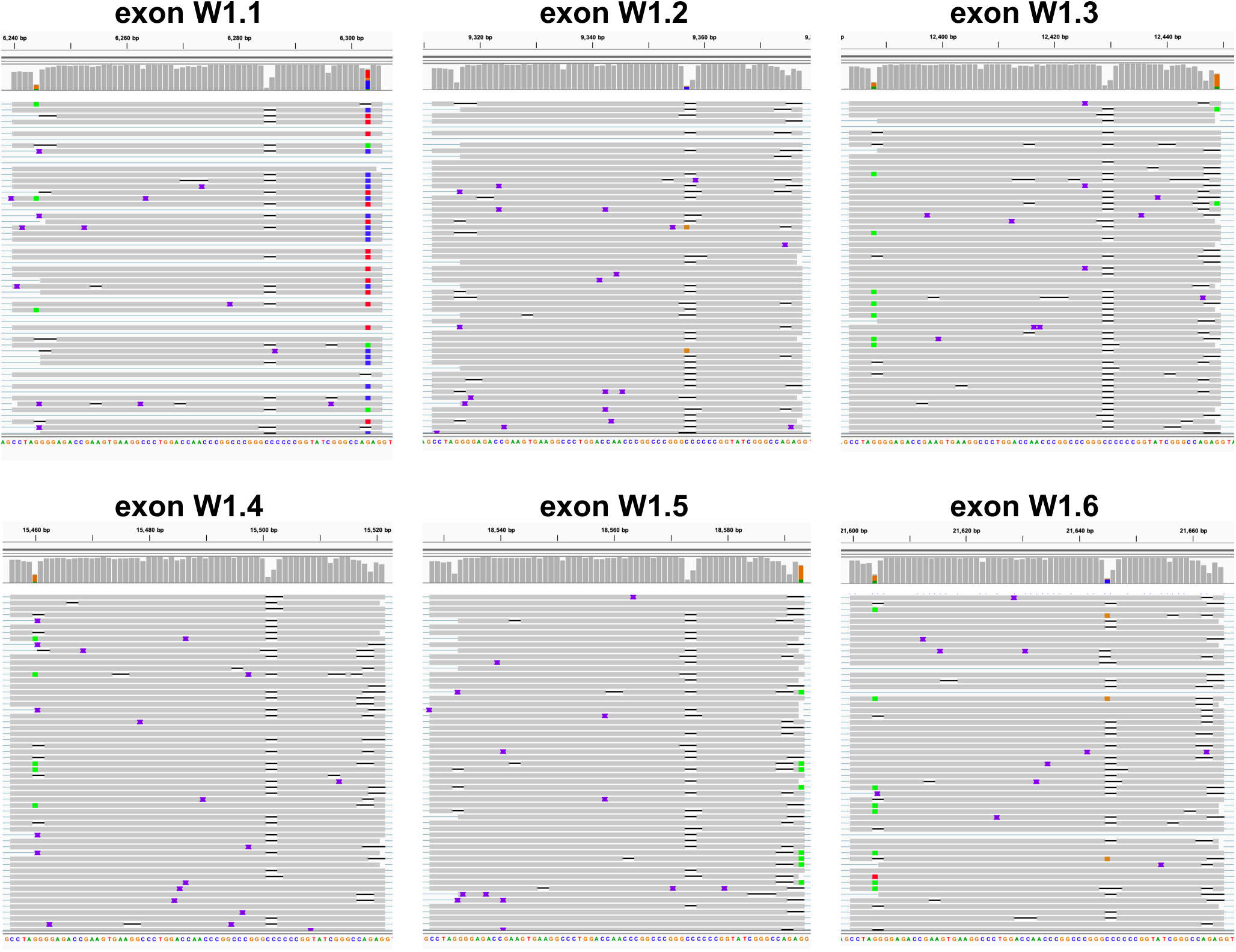
The STOP codon in the W1 exon of WT^HB9^ is in the first exon. Visualisations (using Integrated Genome Browser) of the BAM file of the WT^HB9^-LCL reads aligned to the WT^w^ genome, showing each of the six W1 exons, with genome positions (genome position 1 is the Mlu I restriction site in oriP) marked above. Mismatches of reads with consensus sequence are indicated with coloured boxes, and summarised across all reads in the bars at the top. Grey indicates matches to consensus; black line is deletion and blue flanged element insertion, both relative to consensus.

**Supporting Figure S3.**
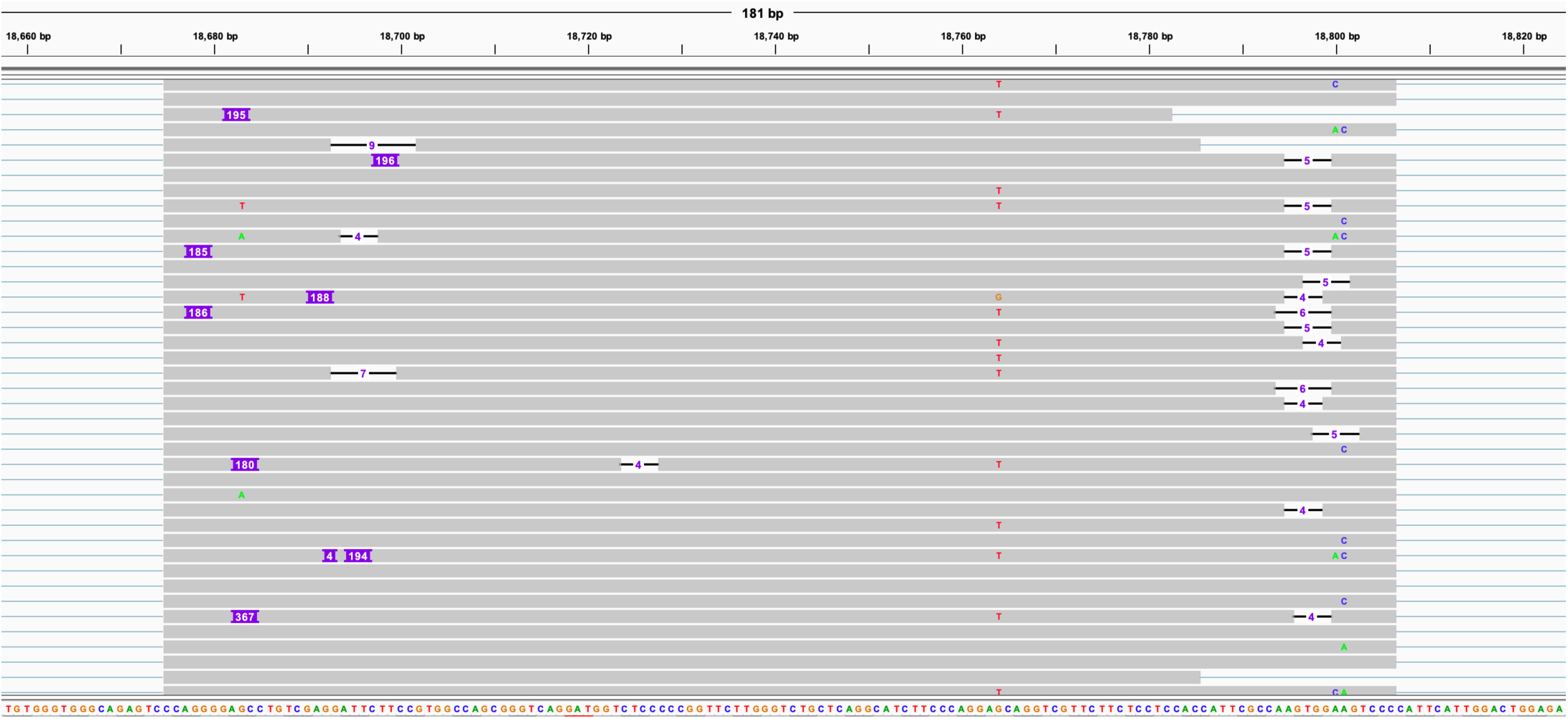
Misalignment of reads to IR1 is evident as multiple exon pair insertions. Visualisation, using integrated genome browser (indels <4 nt are hidden), of reads aligned to a W2 exon. Un-aligned insertions are indicated by blue bars containing white writing. Expected size of W1-W2 exon pair is 193 nt (W1-W2Δ is 172nt).

**Supporting Figure S4.**
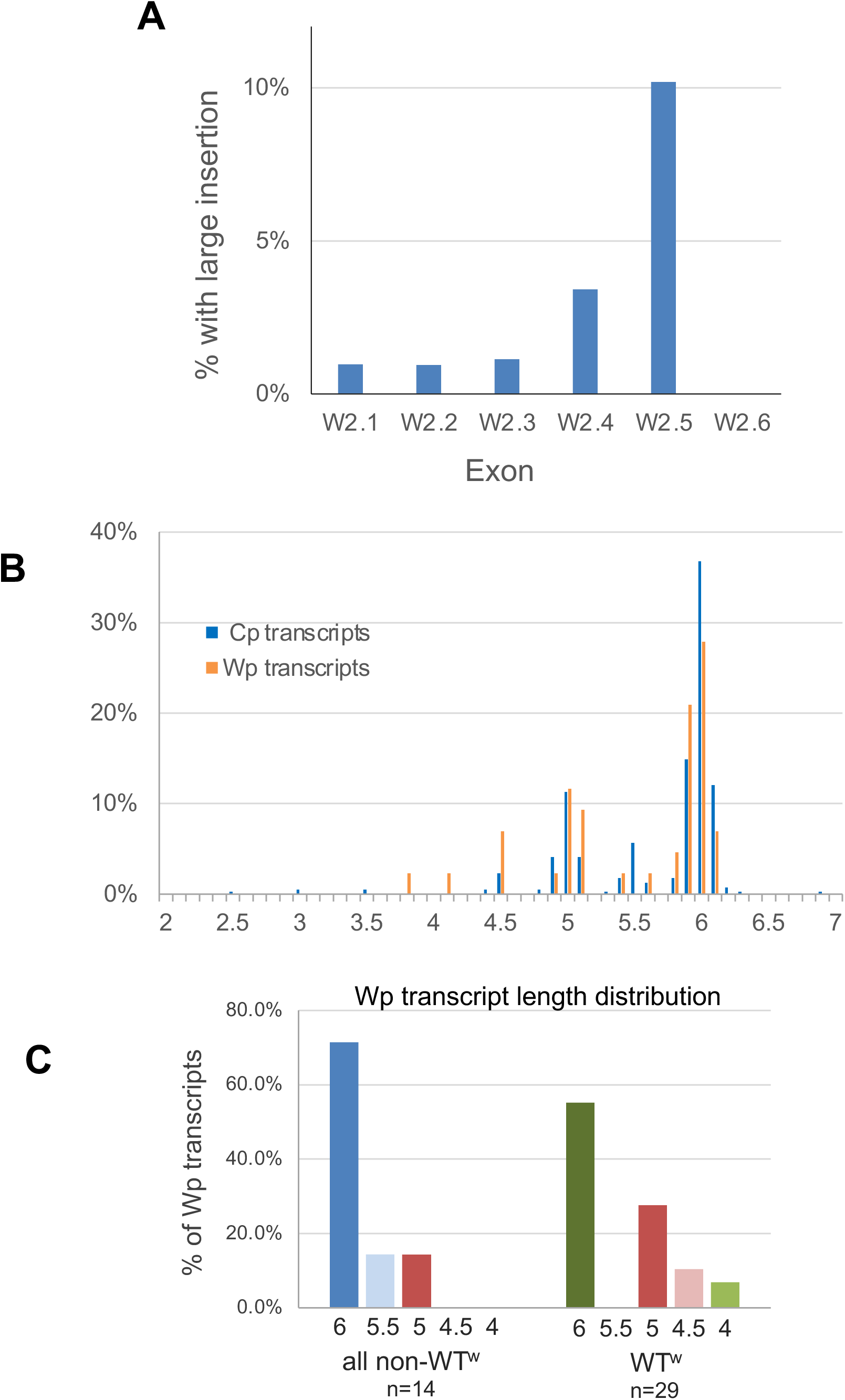
Validation of IR1 counting strategy. **(A)** Frequency of mis-mapping in different W2 exons in IR1 (repeat number is after the “W2.” Percent with insertion is defined as having a number of nucleotides mapped to the exon that is over 1.5 times the ≈130 nucleotides normally observed aligning to that exon. **(B)** The percentage Cp or Wp transcripts (that also map to at least one canonical exon downstream of IR1) with calculated IR1 repeat numbers, plotted as 0.1 repeat bins (centred on integer numbers). The read numbers cluster around 0.5 repeat values, suggesting this approach is effective at estimating the number of IR1 exons in each transcript. **(C**) The distribution of IR1 length estimates of Wp reads, comparing WT^w^ with the other three samples.

## Notes

https://www.ebi.ac.uk/ena/browser/view/PRJEB83447

https://ebv.org.uk

### Competing Interest Statement

The authors have declared no competing interest.

### Summary of Updates

Various clarifications and renaming of some viruses for clarity. Improvement of statistical analysis and inclusion of annotation files for prototype EBV based on our observations.

## References

1. Delecluse HJ, Hilsendegen T, Pich D, Zeidler R, Hammerschmidt W. Propagation and recovery of intact, infectious Epstein-Barr virus from prokaryotic to human cells. Proc Natl Acad Sci U S A 1998;95(14):8245–8250.

2. O’Grady T, Wang X, Höner Zu Bentrup K, Baddoo M, Concha M et al. Global transcript structure resolution of high gene density genomes through multi-platform data integration. Nucleic Acids Research 2016;44(18):e145–e145.

3. Szymula A, Palermo RD, Bayoumy A, Groves IJ, Ba Abdullah M et al. Epstein-Barr virus nuclear antigen EBNA-LP is essential for transforming naïve B cells, and facilitates recruitment of transcription factors to the viral genome. PLOS Pathogens 2018;14(2):e1006890.

4. Cable JM, Wongwiwat W, Grabowski JC, White RE, Luftig MA. Sp140L Is a Novel Herpesvirus Restriction Factor. bioRxiv 2024:2024.2012.2013.628399.

5. Kempkes B, Ling PD. EBNA2 and Its Coactivator EBNA-LP. Curr Top Microbiol Immunol 2015;391:35–59.

6. Styles CT, Paschos K, White RE, Farrell PJ. The Cooperative Functions of the EBNA3 Proteins Are Central to EBV Persistence and Latency. *Pathogens (Basel, Switzerland)*, Review 2018;7(1):31.

7. Altmann M, Pich D, Ruiss R, Wang J, Sugden B et al. Transcriptional activation by EBV nuclear antigen 1 is essential for the expression of EBV’s transforming genes. Proceedings of the National Academy of Sciences 2006;103(38):14188–14193.

8. Yoo L, Speck SH. Determining the role of the Epstein-Barr virus Cp EBNA2-dependent enhancer during the establishment of latency by using mutant and wild-type viruses recovered from cottontop marmoset lymphoblastoid cell lines. J Virol 2000;74(23):11115–11120.

9. Yoo LI, Woloszynek J, Templeton S, Speck SH. Deletion of Epstein-Barr virus regulatory sequences upstream of the EBNA gene promoter Wp1 is unfavorable for B-Cell immortalization. J Virol 2002;76(22):11763–11769.

10. Austin PJ, Flemington E, Yandava CN, Strominger JL, Speck SH. Complex transcription of the Epstein-Barr virus BamHI fragment H rightward open reading frame 1 (BHRF1) in latently and lytically infected B lymphocytes. Proc Natl Acad Sci U S A 1988;85(11):3678–3682.

11. Rogers RP, Woisetschlaeger M, Speck SH. Alternative splicing dictates translational start in Epstein-Barr virus transcripts. *The EMBO Journal*, Comparative Study 1990;9(7):2273–2277.

12. Cai X, Schäfer A, Lu S, Bilello JP, Desrosiers RC et al. Epstein-Barr virus microRNAs are evolutionarily conserved and differentially expressed. PLOS Pathogens 2006;2(3):e23.

13. Pfeffer S, Zavolan M, Grässer FA, Chien M, Russo JJ et al. Identification of virus-encoded microRNAs. *Science (New York*, NY) 2004;304(5671):734–736.

14. Moss WN, Steitz JA. Genome-wide analyses of Epstein-Barr virus reveal conserved RNA structures and a novel stable intronic sequence RNA. BMC Genomics 2013;14:543.

15. Xing L, Kieff E. Epstein-Barr virus BHRF1 micro- and stable RNAs during latency III and after induction of replication. The Journal of Virology 2007;81(18):9967–9975.

16. Rosa MD, Gottlieb E, Lerner MR, Steitz JA. Striking similarities are exhibited by two small Epstein-Barr virus-encoded ribonucleic acids and the adenovirus-associated ribonucleic acids VAI and VAII. Mol Cell Biol 1981;1(9):785–796.

17. Howe JG, Shu MD. Epstein-Barr virus small RNA (EBER) genes: unique transcription units that combine RNA polymerase II and III promoter elements. Cell 1989;57(5):825–834.

18. Laux G, Economou A, Farrell PJ. The terminal protein gene 2 of Epstein-Barr virus is transcribed from a bidirectional latent promoter region. J Gen Virol 1989;70 (Pt 11):3079–3084.

19. Chen H-S, Martin KA, Lu F, Lupey LN, Mueller JM et al. Epigenetic deregulation of the LMP1/LMP2 locus of Epstein-Barr virus by mutation of a single CTCF-cohesin binding site. Journal of virology 2014;88(3):1703–1713.

20. Smith PR, de Jesus O, Turner D, Hollyoake M, Karstegl CE et al. Structure and coding content of CST (BART) family RNAs of Epstein-Barr virus. The Journal of Virology 2000;74(7):3082–3092.

21. Al-Mozaini M, Bodelon G, Karstegl CE, Jin B, Al-Ahdal M et al. Epstein-Barr virus BART gene expression. J Gen Virol 2009;90(Pt 2):307–316.

22. Marquitz AR, Raab-Traub N. Host Gene Expression Is Regulated by Two Types of Noncoding RNAs Transcribed from the Epstein-Barr Virus BamHI A Rightward Transcript Region. Journal of virology 2015;89(22):11256–11268.

23. Edwards RH, Marquitz AR, Raab-Traub N. Epstein-Barr virus BART microRNAs are produced from a large intron prior to splicing. Journal of virology 2008;82(18):9094–9106.

24. Marquitz AR, Raab-Traub N. The role of miRNAs and EBV BARTs in NPC. Semin Cancer Biol 2012;22(2):166–172.

25. Iizasa H, Kim H, Kartika AV, Kanehiro Y, Yoshiyama H. Role of Viral and Host microRNAs in Immune Regulation of Epstein-Barr Virus-Associated Diseases. Front Immunol 2020;11:367.

26. Fachko DN, Goff B, Chen Y, Skalsky RL. Functional Targets for Epstein-Barr Virus BART MicroRNAs in B Cell Lymphomas. Cancers (Basel) 2024;16(20).

27. Jain M, Abu-Shumays R, Olsen HE, Akeson M. Advances in nanopore direct RNA sequencing. Nat Methods 2022;19(10):1160–1164.

28. Workman RE, Tang AD, Tang PS, Jain M, Tyson JR et al. Nanopore native RNA sequencing of a human poly(A) transcriptome. *Nature Methods*, OriginalPaper 2019;16(12):1297–1305.

29. Ba abdullah MM, Palermo RD, Palser AL, Grayson NE, Kellam P et al. Heterogeneity of the Epstein-Barr Virus (EBV) Major Internal Repeat Reveals Evolutionary Mechanisms of EBV and a Functional Defect in the Prototype EBV Strain B95-8. Journal of virology 2017;91(23):e00920–00917.

30. Poling BC, Price AM, Luftig MA, Cullen BR. The Epstein-Barr virus miR-BHRF1 microRNAs regulate viral gene expression in cis. Virology 2017;512:113–123.

31. Allan GJ. Structure and function of the Epstein-Barr virus leader protein. Doctor of Philosophy Thesis, University of London; 1991.

32. Majerciak V, Yang W, Zheng J, Zhu J, Zheng Z-M. A genome-wide Epstein-Barr virus polyadenylation map and its antisense RNA to EBNA. Journal of virology 2018:JVI.01593–01518.

33. Djavadian R, Hayes M, Johannsen E. CAGE-seq analysis of Epstein-Barr virus lytic gene transcription: 3 kinetic classes from 2 mechanisms. PLOS Pathogens 2018;14(6):e1007114.

34. Concha M, Wang X, Cao S, Baddoo M, Fewell C et al. Identification of new viral genes and transcript isoforms during Epstein-Barr virus reactivation using RNA-Seq. J Virol 2012;86(3):1458–1467.

35. Speck SH, Pfitzner A, Strominger JL. An Epstein-Barr virus transcript from a latently infected, growth-transformed B-cell line encodes a highly repetitive polypeptide. Proceedings of the National Academy of Sciences 1986;83(24):9298–9302.

36. Sample J, Brooks L, Sample C, Young L, Rowe M et al. Restricted Epstein-Barr virus protein expression in Burkitt lymphoma is due to a different Epstein-Barr nuclear antigen 1 transcriptional initiation site. Proc Natl Acad Sci U S A 1991;88(14):6343–6347.

37. Sadler RH, Raab-Traub N. Structural analyses of the Epstein-Barr virus BamHI A transcripts. J Virol 1995;69(2):1132–1141.

38. Yamamoto T, Iwatsuki K. Diversity of Epstein-Barr virus BamHI-A rightward transcripts and their expression patterns in the lytic and latent infections. Journal of medical microbiology 2012.

39. Baer R, Bankier AT, Biggin MD, Deininger PL, Farrell PJ et al. DNA sequence and expression of the B95-8 Epstein-Barr virus genome. Nature 1984;310(5974):207–211.

40. Farrell PJ. DNA Viruses: Epstein-Barr virus (EBV). In: O’Brien SJ (editor). Genetic Maps - Locus Maps of Complex Genomes, Book 1: Viruses, 6th Edition: Cold Spring Harbour Laboratory Press; 1993. pp. 120–133.

41. de Jesus O, Smith PR, Spender LC, Elgueta Karstegl C, Niller HH et al. Updated Epstein-Barr virus (EBV) DNA sequence and analysis of a promoter for the BART (CST, BARF0) RNAs of EBV. The Journal of general virology 2003;84(Pt 6):1443–1450.

42. Arvey A, Tempera I, Tsai K, Chen HS, Tikhmyanova N et al. An atlas of the Epstein-Barr virus transcriptome and epigenome reveals host-virus regulatory interactions. Cell Host Microbe 2012;12(2):233–245.

43. Wang Y, Ungerleider N, Hoffman BA, Kara M, Farrell PJ et al. A Polymorphism in the Epstein-Barr Virus EBER2 Noncoding RNA Drives In Vivo Expansion of Latently Infected B Cells. mBio 2022;13(3):e0083622.

44. Fulop A, Torma G, Moldovan N, Szenthe K, Banati F et al. Integrative profiling of Epstein-Barr virus transcriptome using a multiplatform approach. Virol J 2022;19(1):7.

45. Ungerleider N, Concha M, Lin Z, Roberts C, Wang X et al. The Epstein Barr virus circRNAome. PLOS Pathogens 2018;14(8):e1007206.

46. Shekhar R, O’Grady T, Keil N, Feswick A, Amador DAM et al. High-density resolution of the Kaposi’s sarcoma associated herpesvirus transcriptome identifies novel transcript isoforms generated by long-range transcription and alternative splicing. Nucleic Acids Res 2024;52(13):7720–7739.

47. Morgan M, Shiekhattar R, Shilatifard A, Lauberth SM. It’s a DoG-eat-DoG world-altered transcriptional mechanisms drive downstream-of-gene (DoG) transcript production. Mol Cell 2022;82(11):1981–1991.

48. Lasda EL, Blumenthal T. Trans-splicing. Wiley Interdiscip Rev RNA 2011;2(3):417–434.

49. Woisetschlaeger M, Yandava CN, Furmanski LA, Strominger JL, Speck SH. Promoter switching in Epstein-Barr virus during the initial stages of infection of B lymphocytes. Proceedings of the National Academy of Sciences 1990;87(5):1725–1729.

50. Alfieri C, Birkenbach M, Kieff E. Early events in Epstein-Barr virus infection of human B lymphocytes. Virology 1991;181(2):595–608.

51. Allday MJ, Crawford DH, Griffin BE. Epstein-Barr virus latent gene expression during the initiation of B cell immortalization. The Journal of general virology 1989;70 (Pt 7)(7):1755–1764.

52. Kelly GL, Long HM, Stylianou J, Thomas WA, Leese A et al. An Epstein-Barr virus anti-apoptotic protein constitutively expressed in transformed cells and implicated in burkitt lymphomagenesis: the Wp/BHRF1 link. PLOS Pathogens 2009;5(3):e1000341.

53. Xing L, Kieff E. cis-Acting effects on RNA processing and Drosha cleavage prevent Epstein-Barr virus latency III BHRF1 expression. Journal of virology 2011;85(17):8929–8939.

54. Cao S, Moss W, O’Grady T, Concha M, Strong MJ et al. New Noncoding Lytic Transcripts Derived from the Epstein-Barr Virus Latency Origin of Replication, oriP, Are Hyperedited, Bind the Paraspeckle Protein, NONO/p54nrb, and Support Viral Lytic Transcription. J Virol 2015;89(14):7120–7132.

55. Nguyen Quang N, Goudey S, Segeral E, Mohammad A, Lemoine S et al. Dynamic nanopore long-read sequencing analysis of HIV-1 splicing events during the early steps of infection. Retrovirology 2020;17(1):25.

56. Murat P, Zhong J, Lekieffre L, Cowieson NP, Clancy JL et al. G-quadruplexes regulate Epstein-Barr virus-encoded nuclear antigen 1 mRNA translation. Nat Chem Biol 2014;10(5):358–364.

57. Zeglinski K, Montellese C, Ritchie ME, Alhamdoosh M, Vonarburg C et al. An optimised protocol for quality control of gene therapy vectors using Nanopore direct RNA sequencing. bioRxiv 2024:2023.2012.2003.569756.

58. Kwok H, Wu CW, Kellam P, Chiang AKS. Genomic diversity of Epstein-Barr virus genomes isolated from primary nasopharyngeal carcinoma biopsy samples. Journal of virology 2014;88(18):10662–10672.

59. Wang J, Yang L, Cheng A, Tham CY, Tan W et al. Direct RNA sequencing coupled with adaptive sampling enriches RNAs of interest in the transcriptome. Nat Commun 2024;15(1):481.

60. Lee N, Moss WN, Yario TA, Steitz JA. EBV Noncoding RNA Binds Nascent RNA to Drive Host PAX5 to Viral DNA. Cell 2015;160(4):607–618.

61. Naarmann-de Vries IS, Zorbas C, Lemsara A, Piechotta M, Ernst FGM et al. Comprehensive identification of diverse ribosomal RNA modifications by targeted nanopore direct RNA sequencing and JACUSA2. RNA Biol 2023;20(1):652–665.

62. SoRelle ED, Reinoso-Vizcaino NM, Horn GQ, Luftig MA. Epstein-Barr virus perpetuates B cell germinal center dynamics and generation of autoimmune-associated phenotypes in vitro. Front Immunol 2022;13:1001145.

63. SoRelle ED, Dai J, Bonglack EN, Heckenberg EM, Zhou JY et al. Single-cell RNA-seq reveals transcriptomic heterogeneity mediated by host-pathogen dynamics in lymphoblastoid cell lines. Elife 2021;10.

64. White RE, Calderwood MA, Whitehouse A. Generation and precise modification of a herpesvirus saimiri bacterial artificial chromosome demonstrates that the terminal repeats are required for both virus production and episomal persistence. The Journal of general virology 2003;84(Pt 12):3393–3403.

65. Wick RR, Judd LM, Holt KE. Performance of neural network basecalling tools for Oxford Nanopore sequencing. Genome Biol 2019;20(1):129.

66. Li H. New strategies to improve minimap2 alignment accuracy. Bioinformatics 2021;37(23):4572–4574.

67. Li H. Minimap2: pairwise alignment for nucleotide sequences. Bioinformatics 2018;34(18):3094–3100.

68. Loman NJ, Quick J, Simpson JT. A complete bacterial genome assembled de novo using only nanopore sequencing data. Nat Methods 2015;12(8):733–735.

69. Donovan-Banfield I, Turnell AS, Hiscox JA, Leppard KN, Matthews DA. Deep splicing plasticity of the human adenovirus type 5 transcriptome drives virus evolution. Commun Biol 2020;3(1):124.

70. Abebe JS, Alwie Y, Fuhrmann E, Leins J, Mai J et al. Nanopore guided annotation of transcriptome architectures. mSystems 2024;9(7):e0050524.

71. Wade-Martins R, Frampton J, James MR. Long-term stability of large insert genomic DNA episomal shuttle vectors in human cells. Nucleic Acids Research 1999;27(7):1674–1682.

